# Targeting the amino acid metabolism in Lung Sarcopenia: A Systems Engineering Approach

**DOI:** 10.1101/2025.06.30.662260

**Authors:** Gautam Kumar, Shweta Khandibharad, Shailza Singh

## Abstract

Non-small cell lung cancer (NSCLC) is the most prevalent subtype of lung cancer and a leading cause of cancer-related mortality worldwide. Literature evidences indicates a strong association between systemic inflammation, driven by cytokines such as Interleukin-6 (IL-6), and the development of NSCLC-associated sarcopenia. However, the immuno-metabolic underpinnings that link tumor-derived IL-6 signaling to skeletal muscle degradation remain incompletely understood. We developed a comprehensive immuno-metabolic mathematical model to investigate how IL-6 signaling influences branched-chain amino acids (BCAA) metabolism and redox homeostasis in the context of NSCLC-induced sarcopenia by understanding two key causes of sarcopenia which are malnutrition and redox homeostasis. Our model integrates IL-6-mediated activation of the Janus Kinase (JAK)/Signal transducer and activator of transcription 3 (STAT3) pathway, highlighting the differential roles of STAT3 phosphorylation at S727 (mitochondrial electron transport chain (ETC) activation) and Y705 (nuclear translocation and transcriptional regulation of oncogenes Hypoxia-Inducible Factor 1, alpha (HIF-1α) and cellular Myc (c-Myc). These transcription factors reprogram tumor metabolism by upregulating glucose and amino acid transporters, including those specific to BCAAs. In skeletal muscle, elevated IL-6 disrupts insulin signaling via Suppressor of cytokine signaling 3 (SOCS3), suppresses Mechanistic Target of Rapamycin Complex 1 (mTORC1) activation, and enhances oxidative stress, collectively contributing to impaired protein synthesis, increased proteolysis, and muscle wasting. BCAA dysregulation further impairs metabolic balance and exacerbates sarcopenia through defective Tricarboxylic Acid Cycle (TCA) input and epigenetic modulation via acetyl coenzyme A (acetyl-CoA) and S-Adenosylmethionine (SAM). Our model reveals a central role for IL-6–driven metabolic rewiring, particularly BCAA utilization and redox imbalance, in promoting NSCLC-induced sarcopenia. These findings underscore the dual impact of IL-6 on tumor progression and systemic muscle degradation, and provide a framework for evaluating therapeutic strategies that target IL-6/STAT3 signaling and amino acid metabolism via STAT3 and acetyl-CoA cross talk to mitigate NSCLC-induced sarcopenia.

**Authors Summary:** Lung cancer is the leading cause of cancer-related deaths worldwide, and NSCLC accounts for approximately 85% of all cases. Beyond its direct effects, NSCLC can induce systemic conditions such as sarcopenia which is a condition leading to progressive loss of skeletal muscle mass and function through chronic inflammation. A key player in this process is IL-6, a cytokine overexpressed and secreted by tumor cells and cells in the tumor microenvironment. In this study, we developed a systems-level mathematical model to explore how IL-6 contributes to NSCLC-induced sarcopenia by disrupting the metabolism of BCAAs, which are essential for muscle protein synthesis and energy homeostasis. Our model shows that IL-6 activates the JAK/STAT3 signaling pathway in cancer cells, promoting tumor survival and metabolic reprogramming which may simultaneously impair nutritional uptake in muscle cells. IL-6 expressed by Tumor micro environment (TME) may disrupt BCAA uptake and utilization, inhibit anabolic signaling, and increase oxidative stress in muscle tissue, thereby accelerating muscle atrophy. By highlighting the interlinked roles of IL-6 signaling and BCAA metabolism in tumor progression and metabolic reprogramming by cancer cells, our findings offer new insights into the metabolic basis of NSCLC-induced sarcopenia and potential intervention points for therapy.

## Introduction

Lung cancer is one of the most commonly diagnosed cancers and the leading cause of cancer-related deaths worldwide [1]. According to the estimates from the GLOBOCAN 2022, more than 2.48 million new cases of lung cancer (12.4 %) and 1.82 million deaths from lung cancer (18.7 %) occurred in 2022, which present lung cancer as the most common type of cancer and the first leading cause of cancer mortalities [2]. Lung cancers are broadly classified into two types; Small cell lung cancers (SCLC) and NSCLC [3]. SCLC is one of the most aggressive and rapidly growing lung cancers, comprising almost 15% of all lung cancers. SCLC is frequently detected in its advanced stages and spreads quickly to numerous locations [4]. NSCLC is the most common type of lung cancer and accounts for about 85% of all lung cancers. NSCLC can be divided into three main types; Adenocarcinoma, Squamous cell carcinoma and large cell carcinoma [5]. Adenocarcinoma is found in the gland of the lung that produces mucus and is the most common type of NSCLC. Adenocarcinomas comprise more than 40% of all lung cancers and it arises in the outer, or peripheral, areas of the lung [6]. The Squamous Cell Carcinomas accounts for approximately 25-30% of all lung cancer and it grows commonly in the central areas around main bronchi [7]. Large cell carcinomas are the least common type of NSCLC, accounting for 10% of all lung cancers and have a high tendency to spread to the lymph nodes and distant sites [8]. Surgery is the preferred treatment for non-small cell lung cancer (NSCLC) in its early stages. Radiation therapy is used when surgery is not an option. Systemic therapies like chemotherapy, targeted therapy, and immunotherapy are also available [9]. Platinum-based combination chemotherapy is still the standard treatment for advanced cases, while targeted therapies are prescribed for tumors with specific genetic alterations like in epidermal growth factor receptor (EGFR), and anaplastic lymphoma kinase (ALK) [10]. Immune Checkpoint inhibitors, including pembrolizumab and nivolumab, are now often used in immunotherapy, especially for patients without mutations. Programmed cell death protein 1 (PD-1)/Programmed Death Ligand 1 (PD-L1) axis immune checkpoint inhibition is the first-line treatment for most patients with localized and metastatic NSCLC [11]. Bispecific antibodies and antibody-drug conjugates are examples of novel agents that are being developed for tumors that are resistant or advanced [12].

The TME is a dynamic and complex network of, immune cells, stromal cells, cancer cells, extracellular matrix (ECM), and signaling molecules. It plays important role in tumor development, invasion, metastasis, immune evasion, and treatment resistance [13]. Tumor-associated macrophages (TAMs) and cancer-associated fibroblasts (CAFs) release cytokines and growth factors that inhibit anti-tumor immune responses and encourage tumor growth [14]. Cancer cell invasion and migration are facilitated by the modification of the extracellular matrix. Using components like Vascular Endothelial Growth Factor (VEGF), hypoxic conditions inside tumors cause angiogenesis, which results in the development of aberrant blood vessels that aid in the growth and spread of the tumor [15]. Furthermore, the TME promotes resistance to treatment by creating an immunosuppressive milieu that impairs the effectiveness of treatments like chemotherapy and immune checkpoint inhibitors by limiting immune cell infiltration, upregulating immunosuppressive molecules like cluster of differentiation 73 (CD73), and activating survival pathway [16].

The cells in TME secrets cytokines which are small proteins that regulate immune responses and may play dual roles in cancer progression [17]. Anti-inflammatory cytokines like IL-6 trans signaling, Tumor Growth Factor beta (TGF-β), Interleukin-4 (IL-4), Interleukin-13 (IL-13) and Interleukin-1 beta (IL-1β) contribute to tumor initiation, growth, metastasis, and resistance to treatment by promoting chronic inflammation and immune evasion. Conversely, cytokines such as Interferon alpha (IFN-α), Interferon gamma (IFN-γ), Interleukin-2 (IL-2), Interleukin-12 (IL-12), and Interleukin-15 (IL-15) exhibit antitumor effects by enhancing immune surveillance, activating cytotoxic T cells and natural killer (NK) cells, and inhibiting tumor cell proliferation [18]. Therapeutic strategies targeting cytokines include recombinant cytokine therapies, checkpoint inhibitors, and cytokine antagonists, aiming to modulate the TME and improve treatment efficacy. However, the complex roles of cytokines requires precise optimization and delivery to achieve desired therapeutic outcomes [19].

IL-6 is a multifunctional cytokine that significantly contributes to tumor progression through various mechanisms. It promotes cancer cell proliferation, survival, and invasiveness by activating signaling pathways such as (JAK)/(STAT3) and Nuclear factor kappa-light-chain-enhancer of activated B cells (NF-κB), often establishing autocrine loops that sustain malignant phenotypes [20]. IL-6 also facilitates epithelial-mesenchymal transition (EMT), enhancing metastatic potential, and supports the maintenance of cancer stem cells, contributing to tumor heterogeneity and resistance to therapy [21]. Within the tumor microenvironment, IL-6 modulates immune responses by recruiting immunosuppressive cells like myeloid-derived suppressor cells (MDSCs) and TAMs, thereby diminishing anti-tumor immunity [22]. Additionally, IL-6 induces the expression of vascular endothelial growth factor (VEGF), promoting angiogenesis to supply nutrients and oxygen to the growing tumor [23]. These multifaceted roles of IL-6 in cancer progression underscore its potential as a therapeutic target in oncology.

IL-6 plays an important role in sarcopenia which is marked by progressive decline of muscle mass, strength and its quality through different cellular and molecular mechanisms [24]. In skeletal muscle cells elevated IL-6 levels activate the JAK/STAT3 signaling pathway, leading to the phosphorylation and nuclear translocation of STAT3, which promotes the transcription of genes associated with muscle protein degradation [25]. This process increases the expression of the SOCS3, which not only inhibits further IL-6 signaling but also interferes with insulin signaling by targeting insulin receptor substrate-1 (IRS-1) for degradation. The reduction in Insulin Receptor Substrate-1 (IRS-1) impairs the Phosphatidylinositol 3-kinase catalytic subunit alpha (PI3K)/Ak strain transforming (Akt) pathway, crucial for muscle protein synthesis, thereby exacerbating muscle atrophy [26]. Additionally, IL-6-induced STAT3 activation suppresses anabolic pathways, such as the insulin-like growth factor 1 (IGF-1)/Akt/mTOR axis, further contributing to muscle wasting [27].

In NSCLC induced sarcopenia, IL-6 may activate the JAK/STAT3 signaling pathway and also disrupts BCAA metabolism and dysregulated redox homeostasis in muscle tissue. This pathway suppresses mTORC1 activation and protein synthesis, impairs BCAA utilization, and increases muscle protein breakdown, [28]. while also elevating oxidative stress by boosting reactive oxygen species (ROS) production and reducing antioxidant defenses [29]. These combined effects accelerate muscle wasting even more and create a vicious cycle of inflammation and metabolic imbalance. Therapeutic strategies focus on blocking IL-6 signaling (e.g., with tocilizumab), optimizing BCAA intake, using antioxidants to restore redox balance, and incorporating exercise to counteract muscle loss.

Therapeutic strategies targeting IL-6 signaling pathways are under investigation to mitigate sarcopenia. Pharmacological inhibitors of JAK2 (e.g., AG490) and STAT3 (e.g., C188-9) have demonstrated efficacy in preclinical models by reducing muscle atrophy and preserving muscle mass [30]. Monoclonal antibodies against IL-6 or its receptor, such as tocilizumab, are also being explored for their potential to attenuate inflammation-induced muscle degradation [31]. Complementary approaches, including resistance exercise and nutritional interventions like protein supplementation, remain foundational in managing sarcopenia [32]. These combined strategies aim to restore the balance between muscle protein synthesis and degradation, offering a multifaceted approach to counteract IL-6-mediated muscle loss.

To understand NSCLC induced sarcopenia we have reconstructed immuno-metabolic mathematical model of cancer cell highlighting the roles of IL-6 signaling in metabolic pathways bridged together by BCAA. In the presented model, IL-6 released from tumor cells and TAM promote tumor cell survival and progression by multiple ways. IL-6 when bound to its cell surface receptor Interleukin-6 Receptor Alpha (IL-6Rα) Receptor Tyrosine Kinase (RTK) it starts downstream signaling cascade and phosphorylates and activates JAK1/2 molecules. Now this JAK1/2 molecule further phosphorylates and activates STAT3. When STAT3 is phosphorylated at S727 position it will translocate to mitochondria and bind with complex1 and promote ETC. Additionally, it also shows anti apoptotic effects. But, when IL-6 phosphorylated at Y705 position it dimerizes and translocates inside the nucleus and regulates expression of HIF-1α and c-Myc. The HIF-1α and c-Myc are oncogenes which generally are upregulated in many cancers. It regulates expression of various glucose and amino acid transporters and their enzymes for metabolic reaction. Additionally, STAT3 also promotes metastasis and angiogenesis by expressing N-cadherin, matrix metalloproteases and VEGF. The HIF-1α and c-Myc oncogene upregulate the expression of Glucose transporters (GLUTs) and key glycolytic enzyme; Hexokinase, Pyruvate Kinase Muscle Isozyme Type 2 (PKM2) and Lactate Dehydrogenase A (LDHA) and favour glycolytic pathway more along with Purine and Serine synthesis pathway. Serine is involved in folate cycle and form glycine which along with glutamine form glutathione act as key molecules of redox homeostasis. The folate cycle is coupled with the methionine cycle and it forms SAM, a methyl donor involved in the methylation. Due to overexpression of oncogene, HIF-1α and c-Myc it upregulates the expression of branch chain amino acid (Leucine, Isoleucine and Valine), glutamine, solute carrier family 1 (neutral amino acid transporter), member 5 (SLC1A5) and tryptophan L-type amino acid transporter 1 (LAT1) transporter and allow high molecule influx inside cell. The BCAA molecules after several steps of metabolic reaction it forms acetyl-CoA and Succinyl Coenzyme A (succinyl-CoA). The acetyl-CoA molecules were used as precursor or an intermediate in fatty acid synthesis and participate in histone acetylation. The Succinyl-CoA molecules convert to succinate and enter in the TCA cycle. The aspartate from TCA cycle is the important molecule for synthesis of purine and pyrimidine and it is also involved in urea cycle. Overall IL-6 induced STAT3 molecules are holistically involved in tumor progression and survival by regulating metabolism of glucose, BCAA, tryptophan and glutamine. Redox homeostasis, nucleotide biosynthesis and fatty acid synthesis is also regulated by IL-6. Additionally, these metabolic pathways associated with IL-6 promote metastasis, angiogenesis and epigenetic modification (acetylation and methylation) and inhibits apoptosis; all prominent hallmarks.

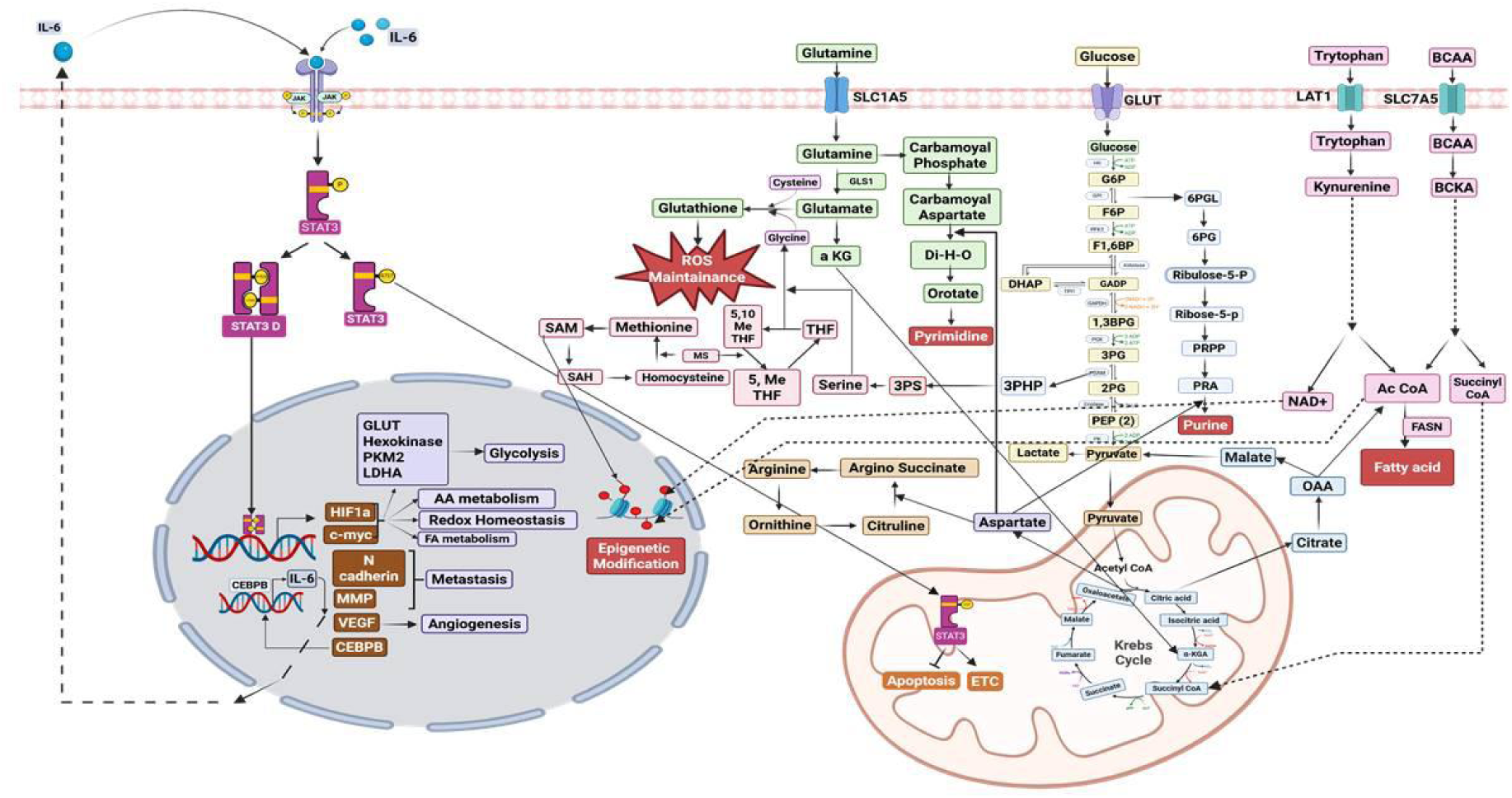

Graphical abstract of the reconstructed mathematical model of NSCLC induced sarcopenia

## Methodology

### Reconstruction and simulation of the NSCLC induced sarcopenia cancer model

To elucidate the mechanistic interplay between NSCLC and sarcopenia, we developed a dynamic, multi-compartmental mathematical model integrating the immuno-metabolic implications of IL-6 signaling with BCAA metabolism. The model features rate laws, quantitative kinetic parameters and initial component concentrations to decipher the implications of nutrition and redox potential on metabolism in NSCLC induced sarcopenia co-morbid model. The model was reconstructed using Cell Designer where immune signaling and metabolic pathways were defined in the form of reactions.

The kinetic rate laws included, Law of Mass action kinetics (for association, dissociation, transcription, translation, and activation reactions), Michaelis-Menten kinetics (for enzyme-catalyzed reactions) and Hill’s kinetics (for transcription reactions).

Initial concentrations of components were derived from literature, which suggests that a cell secretes 10^3^-10^6^ signaling molecules [33]. Using the SBML ODE Solver Library (SOSlib, v1.7.0) solver, the model was simulated for 100 s to obtain concentration versus time graph.

### Principal component analysis of the NSCLC induced sarcopenia cancer model

PCA was performed to reduce dimensionality and identify key components driving sarcopenia progression in NSCLC. The analysis utilized FactoMineR (v4.1.3) and Factoextra (v4.1.3) packages in RStudio (v4.1.1). The input matrix included component concentrations from metabolites and pathway intermediates which were a part of only one signaling pathway versus multiple signaling pathways. PCA helped distinguish and highlight principal components (PCs) with significant contributions having metabolic and inflammatory signatures which may affect sarcopenia induced by NSCLC highlighting.

### Flux Analysis

Flux analysis, a constraint-based method, was employed to assess metabolic flow distributions in the NSCLC induced sarcopenia model under steady-state conditions. The analysis was conducted using COPASI (v4.36.260), a biochemical network simulator, which solves ordinary differential equations (ODEs) to determine reaction fluxes. Briefly, the model was exported in SBML format and was imported on COPASI. In tasks, steady state option was selected. Numerical Resolution was set to 1×10⁻⁹ to ensure precision in flux calculations, Derivation Factor was fixed at 0.001 for stable gradient estimation during optimization, Jacobian & Stability Analysis was Enabled to assess system robustness and dynamic behavior near steady state, Forward/Backward Integration was set to maximum durations of 1×10⁹ (forward) and 1×10⁶ (backward) time units to assure convergence, Newton Iteration Method was activated with an iteration limit of 50 to balance accuracy and computational efficiency. Convergence Criterion was defined by distance and rate thresholds to confirm steady-state attainment. High-flux reactions, critical for network functionality, were identified by setting a threshold of 1000 mol/s.

### Cross talk point

A crosstalk point refers to any direct or indirect interaction between signaling components of two distinct pathways that influence a biological response. These nodes are critical regulators within signaling networks and can significantly impact the overall network dynamics, especially in reconstructed models. This analytical approach helps identify key regulatory elements by examining the network architecture, with a higher crosstalk score indicating a greater potential of a given component to drive inflammatory responses. To pinpoint regulatory nodes, the maximum degree of each node within individual pathways was subtracted from the total degree of the corresponding node in the entire network. Nodes with non-zero values from this analysis are considered crosstalk points within the signaling network.

### Network Construction and analysis

The mathematical model was constructed as an immuno-metabolic undirected network and analyzed on Cytoscape (v3.9.1). The reactants in the model were defined as source and products were defined as targets. The table was imported on cytoscape and layout for visualization was set to circular layout. The model was analysed by choosing the analyze option from the tools tab. Global Network Properties like number of nodes and edges, average number of neighbors, network diameter, network radius, characteristic path length, Clustering coefficient, Network density, Network heterogeneity, network centralization and connected components were evaluated.

### CytoHubba analysis and frequency of occurrence

CytoHubba is a Cytoscape plugin which was used to identify key nodes within the constructed immuno-metabolic networks. Topological analyses were performed using twelve distinct algorithms, including degree, betweenness, closeness, eccentricity, radiality, stress, bottleneck, maximum neighborhood component (MNC), density of maximum neighborhood component (DMNC), edge percolated component (EPC), and maximal clique centrality (MCC). These measures collectively rank nodes based on their centrality and connectivity within the network. The frequency of occurrence of a species in the rank list of all the above-mentioned parameters were highlighted thereby identifying potential regulatory hubs that may play pivotal roles in modulating the immune-metabolic response in NSCLC induced sarcopenia.

### BinGO analysis

The BiNGO (Biological Networks Gene Ontology) Cytoscape plugin was used to functionally annotate the network components. This tool determined overrepresented Gene Ontology (GO) categories within the network’s subgraphs, comparing input components against a reference set. GO enrichment was assessed using a hypergeometric test, and significance was adjusted with the Benjamini-Hochberg correction to control the false discovery rate. The annotation reference genome was *Homo sapiens*. Functional categorization revealed biological processes and molecular functions significantly associated with immuno-metabolism in NSCLC induced sarcopenia

### Cell lines and treatment conditions

A549, H1975 and H1299 cells were cultured in respective medias HF-10 (Ham’s F-10), and RPMI-1640 (Roswell Park Memorial Institute 1640). Total 10^6^ cells were induced with 1 μg Human rIL-6 for 24 h at 37 °C 5% CO_2_. These samples were processed for Western blot, qRT-PCR and the supernatant was used for ELISA.

### Western blot

Protein samples were prepared using RIPA lysis buffer (50 mM Tris-HCl pH 7.4, 150 mM NaCl, 0.25 mM EDTA pH 8.0, 5 mM NaF, 1% Triton X-100, 1% sodium deoxycholate, 1 mM Na3VO4) supplemented with 1X protease inhibitor. Total protein concentration was quantified using the bicinchoninic acid (BCA) assay. For immunoblotting, 20 μg of protein per sample was resolved by SDS-PAGE on 10% polyacrylamide gels under reducing conditions.

Following electrophoresis, proteins were transferred to the PVDF membrane (MDI, SCNX8701) using standard wet transfer protocols. Membranes were blocked with 3% bovine serum albumin (BSA; HiMedia, 23228) in Tris-buffered saline containing 0.1% Tween-20 (TBST) for 1 hour at room temperature with gentle agitation.

Primary antibody incubation was performed overnight at 4°C with GAPDH, (Cell signaling technology, 2118S) P-Y705-STAT3, (Cell signaling technology, 9131S) AKT (Cell signaling technology 9272S), P-AKT (S473) (Cell signaling technology 4060S) and Arg1(Cell signaling technology 93668) (1:1000 dilution in blocking buffer). After three 10-minute TBST washes, membranes were incubated with horseradish peroxidase (HRP)-conjugated goat anti-rabbit IgG secondary antibody (Sigma-Aldrich, A9169; 1:5000 dilution) for 1 hour at room temperature. Following three additional TBST washes, protein bands were visualized using enhanced chemiluminescence (Luminol-Enhancer Solution and H2O2; Cyanagen, XLS070L/XLS070P) and imaged on an Amersham ImageQuant 500 digital imaging system. Densitometric analysis was performed using ImageJ.

### Laser scanning confocal microscopy

Total 10^4^ cells of A549, H1975 and H1299 cell line were seeded in 96 well coverslip bottom plate and incubated for 24 h at 37 °C 5% CO_2_. Post incubation, media was discarded and the cells were washed thrice with 1X PBS. Cells were induced with 20 ng of Human rIL-6 for 24 h at 37 °C 5% CO_2_. Post incubation, the cells received three cycles of washing with 1XPBS using 0.05% Tween 20 (3601181001730 Genei). The cells were then washed with 1XPBS after being fixed with 4% paraformaldehyde (PFA) for 10 minutes at room temperature. Cells were treated with 1X PBST (0.1% TritonX (9002-93-1 ThermoFischer Scientific) in 1X PBS) for 10 minutes at room temperature and later blocked with 3% Bovine Serum Albumin (BSA) (9048-46-8, Sigma-Aldrich) made in 1X PBST for 30 minutes at room temperature. After removing the blocking media from the cells, they were washed three times with 1X PBST. AKT, P-AKT (S473), IL-6, and P-Y705-STAT3 primary antibodies (1:1000) were added to the samples, and the cells were then incubated for two hours at room temperature. They were then washed three times with 1X PBST. The cells were incubated with anti-rabbit IgG (H + L) F(ab′)2 fragment (AlexaFluor 488 Conjugate) ((4412S) Cell Signaling Technology) and mouse phalloidin AlexaFluor 568 ((A12380) ThermoFisher Scientific) (1:500) for 1 hour. They were then washed three times with 1X PBST. After 10 minutes of DAPI counterstaining, cells were washed three times with 1× PBST. Cells were then observed at 60x using an Olympus FV3000 microscope after being washed three times with distilled water. Fiji 1.54f was used to analyze the images.

### Real time-PCR mRNA isolation

To the samples, 500 μL of TRIzol reagent was added followed by 100 μL Chloroform. The 1.5 mL tube was inverted and the samples were gently mixed. For 15 minutes, the samples were incubated at room temperature. Samples were then centrifuged for 30 minutes at 4°C at 12,000 rpm. A fresh 1.5 mL tube was used to collect the top aqueous transparent layer. 500 µL of isopropanol was added to the aqueous layer, and the tubes were gently mixed by inverting them. For ten minutes, the tubes were incubated at room temperature. Following incubation, the samples were centrifuged for 15 minutes at 4°C at 12,000 rpm. The acquired pellet was centrifuged at 8000 rpm for 5 minutes at 4°C, and the supernatant was discarded. The pellet was then twice washed with 75% chilled ethanol. After being allowed to air dry, the pellet was reconstituted in 20 µL of diethylpyrocarbonate (DEPC) water. Nanodrop was used to quantify the extracted RNA and record the 260/280 and 260/230 ratios.

### cDNA synthesis

cDNA was synthesized in 100 µL tubes using 2 µg of mRNA from each sample. We prepared 20µL of cDNA using 0.8 µL of 25X dNTP, 2 µL of 10X RT Primer, 2 µL of 10X RT buffer, 0.5 µL of Reverse Transcriptase, and 2 µg of mRNA according to Nanodrop quantification. The total volume was adjusted using DEPC water. Using a thermal cycler, the samples were heated to 25°C for 10 minutes, then to 37°C for 2 hours, then to 85°C for 5 minutes, and finally to 4°C forever. qRT-PCR

Expression profile for Lactate dehydrogenase (LDHA) (HS01378790_G1 ThermoFischer Scientific), acetyl-CoA carboxylase (ACAC) (HS01046047_M1 ThermoFischer Scientific) and Pyruvate dehydrogenase E1 subunit alpha 1 (Pdha1) (HS01049345_G1 ThermoFischer Scientific) using TaqMan-FAM-MGB chemistry. Differential mRNA expression analysis was performed using a total of 2 µg of cDNA from each sample. 10µL of the reaction mixture was utilized. 5 µL of (2X) TaqMan Fast Universal PCR Master Mix (4366072 ThermoFischer Scientific), 0.5 µL of the Taqman-FAM-MGB probes that are listed above, and 1 µL of cDNA from samples were the chemicals that were added. Nuclease-free water was used to make up the entire volume. The StepOne plus Real Time PCR machine and StepOne software version 2.3 were used to perform the qRT-PCR. The following steps were part of the PCR cycle: holding at 50°C for two minutes, holding at 95°C for twenty seconds, cycling at 95°C for three seconds, and cycling at 60°C for thirty seconds. There were forty cycles in total. For each probe and sample Comparative CT (ΔΔ CT) was recorded. The housekeeping gene used was Actin B (HS01060665-G1 ThermoFischer Scientific). Expression in terms of fold change was recorded for all probes.

### Enzyme-linked immunosorbent assay (ELISA)

Supernatant from each sample was collected and examined for presence of IL-6 using IL-6 Human ELISA kit (KHC0061, Thermo Fisher Scientific) according to the manufacturer’s instructions. Briefly, supernatants were mixed with a 1:100 diluted Biotin-conjugated primary antibody. After that, the mixture was incubated for two hours at room temperature on a shaker. The mix was discarded and the wells were washed four times with wash buffer. Following the washes, to each well 1:100 diluted streptavidin-HRP secondary antibody was added and incubated on shaker at RT for an hour. The wash buffer was then used three times to clean the wells. Following the addition of TMB substrate and thirty minutes of dark incubation, stop solution was added to the wells. A multi spectrophotometer was used to acquire the wells at 450 nm, and colorimetric measurements were recorded for every sample. By extrapolating on a standard curve graph, IL-6 levels were quantified.

### Statistical analysis

The quantitative data represented the result from two independent experiments which produced similar results. The reported data shows the mean and standard deviation (SD) of at least two independent experiments. The dataset was analyzed using the two-way ANOVA for groups with two independent variables. Asterisks denoted represents the statistical significance for differences, with a p-value less than 0.05(*), p-value less than 0.01 (**), p-value less than 0.001 (***) and p-value less than 0.0001 (****). Data analysis for all experiments was performed using GraphPad Prism 5 for Windows, which is a program developed by La Jolla, California-based GraphPad Software (www.graphpad.com).

## Results

### Mathematical models and simulation NSCLC induced Sarcopenia cancer model

We have reconstructed a lung cancer immuno-metabolic mathematical model which was composed of four compartments including cell membrane, cytoplasm, mitochondria and nucleus. A total of 229 components and 123 reactions were incorporated to reconstruct the model. Simulation was performed, using SBML ODE Solver Library SOSlib (1.7.0) available in Cell Designer (4.4.2).

In this model, we integrated IL-6 which may be released from NSCLC tumor cells and TAMs which enhance tumor survival and progression through multiple mechanisms. The reconstructed model highlighted the reactions where IL-6 binding to its receptor (IL-6Rα/RTK) activated the JAK1/2-STAT3 pathway. Phosphorylated STAT3 at S727 translocates to mitochondria to promote the electron transport chain and inhibit apoptosis, while phosphorylation at Y705 leads to nuclear translocation, upregulating oncogenes HIF-1α and c-Myc. These oncogenes increase the expression of glucose and amino acid transporters and key metabolic enzymes, favoring glycolysis, purine, and serine synthesis. Serine and glutamine contribute to redox homeostasis via glutathione production, and the folate cycle, linked to the methionine cycle, produces SAM for methylation. Overexpression of HIF-1α and c-Myc also boosts BCAA, glutamine, and tryptophan transport, fueling metabolic pathways that generate acetyl-CoA and succinyl-CoA for fatty acid synthesis, histone acetylation, and the TCA cycle. Aspartate from the TCA cycle supports nucleotide biosynthesis and the urea cycle. These overall, IL-6-driven STAT3 activation orchestrates tumor metabolism, metastasis, angiogenesis, epigenetic modifications, and apoptosis inhibition which are key hallmarks of cancer progression (Figure 1). We observed the concentration of species including fatty acids, acetyl-CoA, succinate, malonyl coenzyme A (malonyl-CoA), glutathione, pyrimidine, Nicotinamide Adenine Dinucleotide (NAD+), and epigenetic modifiers were high upon model simulation which may play critical roles in maintaining the cancer phenotype (Figure 2).

**Figure 1.**
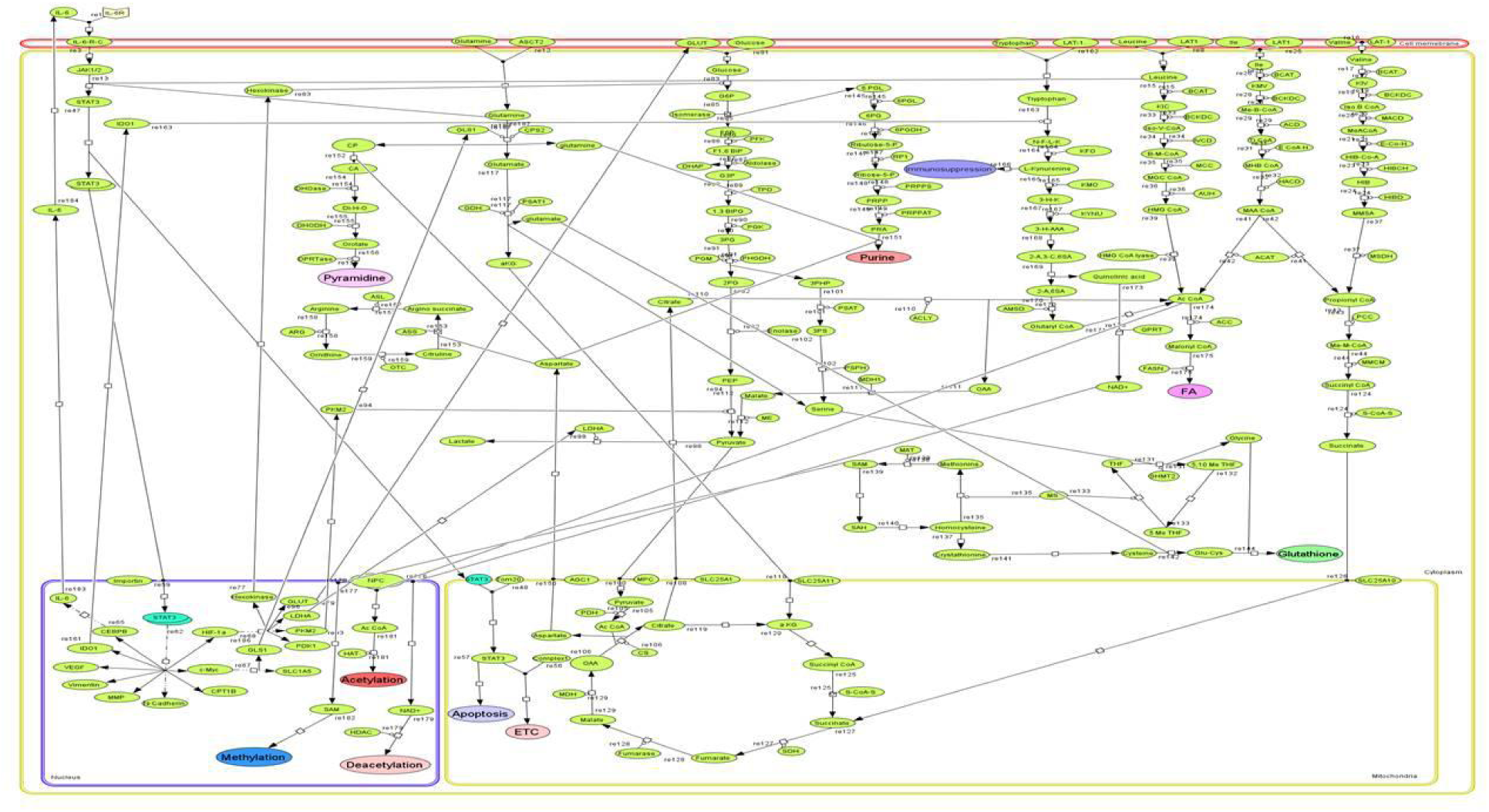
Reconstructed mathematical model of NSCLC inducing sarcopenia through IL-6 signaling and BCAA metabolism

**Figure 2.**
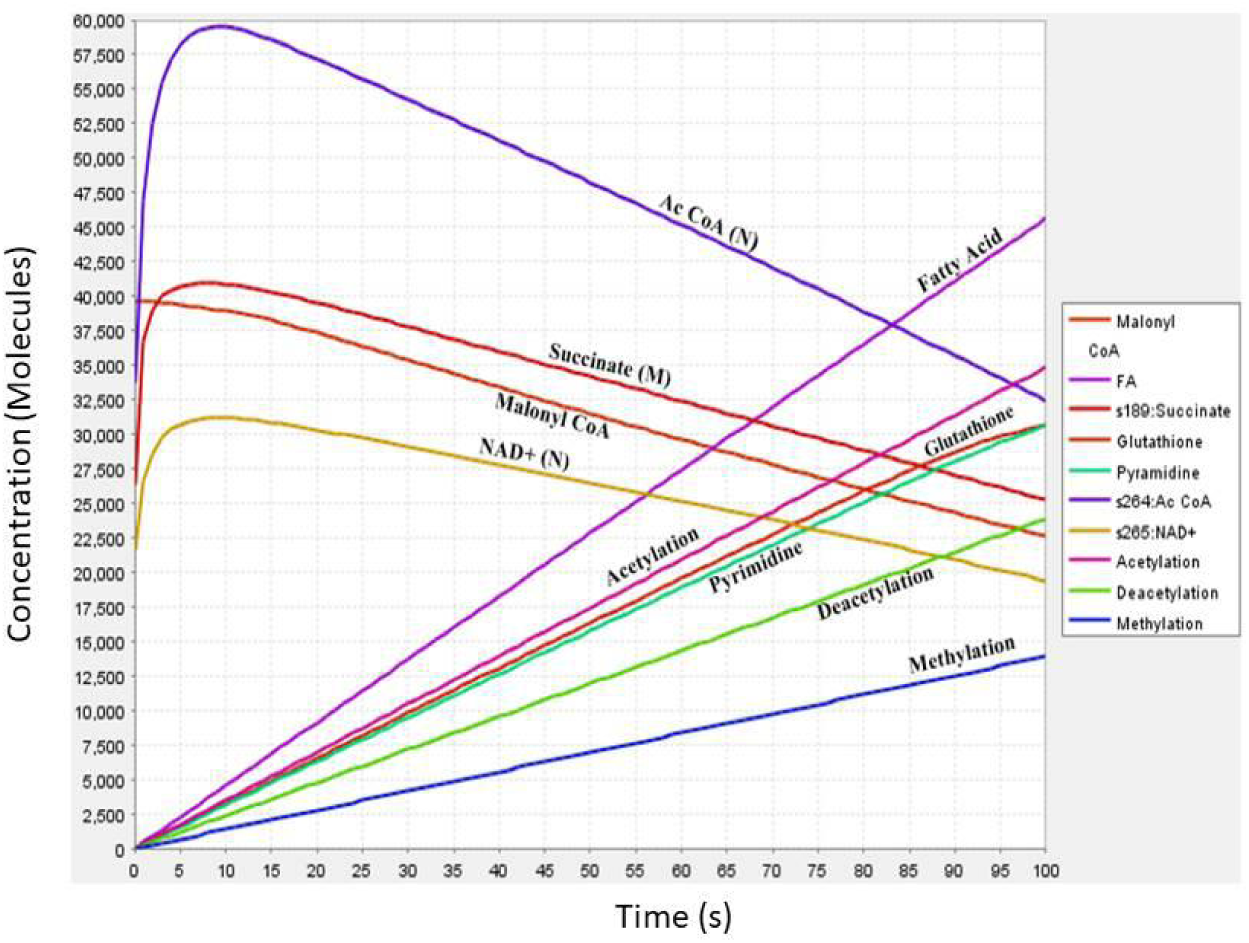
Simulation of the reconstructed mathematical model to obtain concentration versus time plot

### Principal component analysis (PCA)

We had categorized all the species into two groups; one group included species which participated in multiple pathways and the other group comprised species which were involved in single pathway (S1). From PCA analysis the plot covered 50.5% and 49.5% dimension of our data. Furthermore, when we obtained the variable PCA which showed the groups were diverse and had variable PC score of 0.7 (Figure 3a). The individual PCA plot highlighted the principal components which are involved in single, multiple as well as in a unique way balancing both pathways (Figure 3b). Components from multiple pathways included 5,10-Methenyltetrahydrofolate (5,10 methyl THF) from Serine/Folate pathways, acetyl-CoA (M) from Glycolysis/TCA pathways, acetyl-CoA (N) from Acetylation/Leucine/Ile/Glycolysis/Tryptophan pathways and NAD+ from deacetylation/Tryptophan pathways. Fatty acid synthase (FASN) from fatty acid metabolism, malate dehydrogenase (MDH) from TCA cycle, succinyl-Co-A synthetase from TCA cycle, and importin from IL-6 pathway are principal components from individual pathway. PCs which were unique highlighted phosphohydroxy pyruvate (PHP) from Glycolysis/Serine pathways, LDHA from IL-6/Glycolysis pathways, pyruvate dehydrogenase PDH from Glycolysis/TCA pathways, and alpha ketoglutarate (α-KG) from TCA/Glutamine pathways are involved in both pathways (Figure 3c). The individual contribution graph suggests a total of 19 components possessed PC score greater than 1 and are enlisted in the table (Figure 3d and 3e).

**Figure 3.**
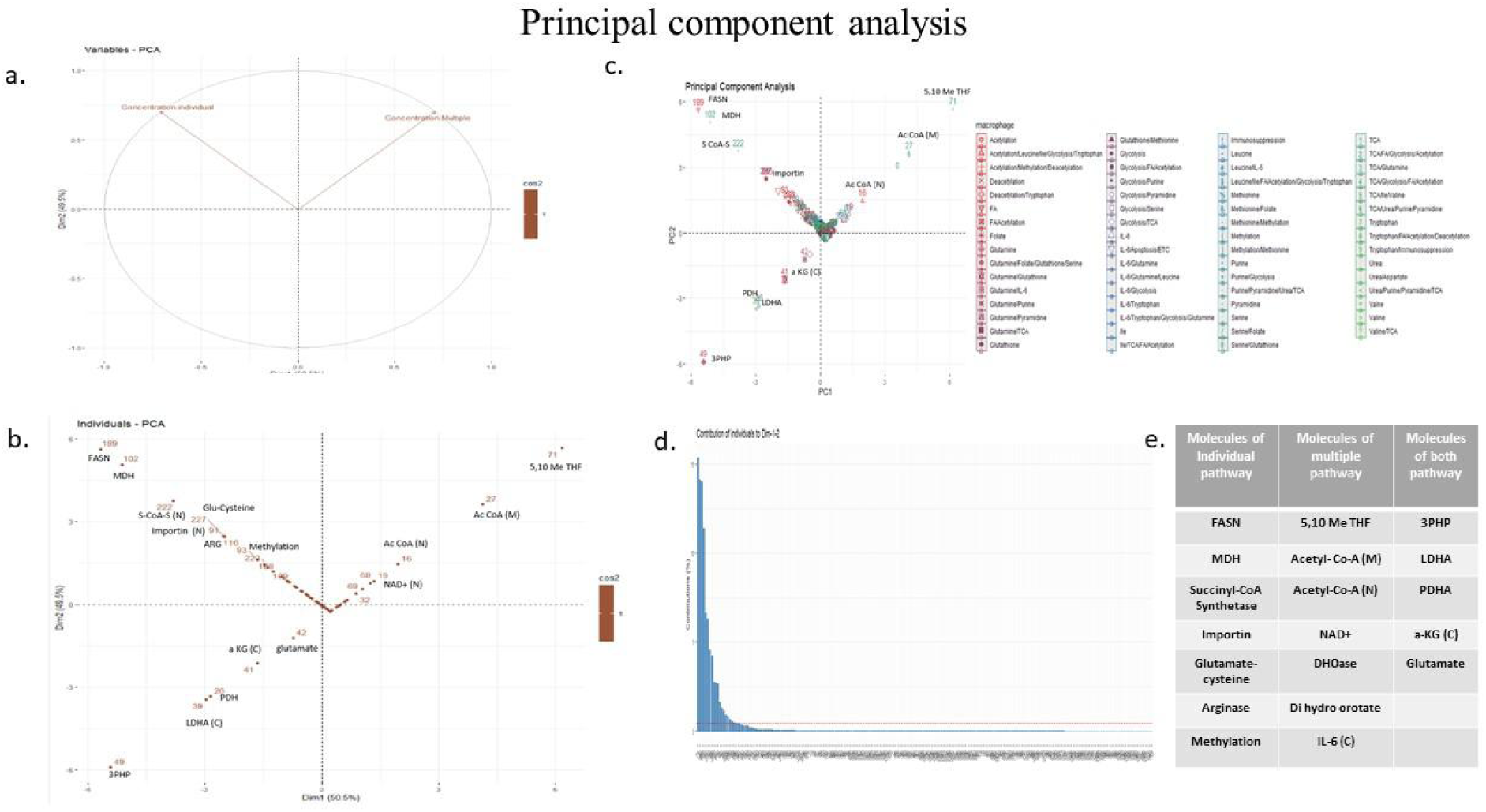
Principal component analysis of the mathematical model- (a) Variable PCA plot (b) Individual PCA plot (c)PCA plot based on pathways (d) Individual contribution plot (e) Table highlighting the components of contribution plot

### Flux analysis

Through comprehensive metabolic flux analysis, found succinate, acetyl-CoA, α-KG, citrate, aspartate, pyruvate, BCAA comprising isoleucine, valine and leucine, epigenetic modifiers (NAD+, SAM) and STAT3 have high flux (Table 1) (S2). These metabolites not only act as intermediates in the TCA cycle and mitochondrial metabolism but also as critical modulators of signaling and gene expression, thereby contributing to the metabolic reprogramming in NSCLC and serve as metabolic markers for muscle wasting characteristic of cancer-induced sarcopenia. Succinate is known role as an oncometabolite in the TME. In NSCLC, high succinate levels induce HIF-1α and promotes glycolytic reprogramming and angiogenesis, fueling tumor growth. In skeletal muscle, this may impair oxidative metabolism, leading to reduced mitochondrial function and enhanced muscle degradation, hallmarks of sarcopenia [34]. In NSCLC cells, elevated acetyl-CoA can sustain anabolic growth by enhancing histone acetylation and promoting the transcription of genes involved in proliferation and metabolic adaptation. In the context of sarcopenia, altered acetyl-CoA dynamics may disrupt muscle-specific gene expression by modifying chromatin accessibility, contributing to atrophy [35] [36]. High flux of citrate and other TCA metabolites suggests enhanced mitochondrial export and cytosolic cleavage to generate acetyl-CoA, linking mitochondrial metabolism to lipid biosynthesis and epigenetic modifications. However, in muscle tissue, diversion of citrate and downstream intermediates away from ATP production may exacerbate energy deficiency, further promoting proteolysis and muscle wasting [37]. Elevated NAD⁺ flux indicates increased activity of redox reactions and deacetylation processes. The deacetylation changes likely facilitate tumor cell adaptation while simultaneously repressing transcriptional programs required for muscle maintenance and regeneration [38] [39].

(Table1).

**Table 1.** High flux reactions from the reconstructed mathematical model.

| Species | Flux mol/sec |
| --- | --- |
| Succinate(C)+SLC25A10-> Succinate(M) | 28449.6 |
| Ac CoA(C)+NPC->Ac CoA (Nu) | 21970.9 |
| NAD+(C) +NPC-> NAD+(Nu) | 8295 |
| SAM(C)+NPC->SAM(Nu) | 6948.9 |
| a KG(C)+SLC25A11-> a KG(M) | 5975.29 |
| Citrate(M)+SLC25A1-> Citrate(C) | 5617.5 |
| Aspartate(M)+AGC1-> Aspartate© | 5617.5 |
| Pyruvate (C)+MPC-> Pyruvate (M) | 4178.8 |
| Ile (PM)+LAT1.-> Ile© | 3200 |
| Valine (PM)+LAT-1-> Valine© | 3200 |
| Leucine (PM)+LAT1->Leucine© | 1600 |
| STAT3 (NM)+Importin->STAT3(Nu) | 1250 |

### Cross talk point analysis

Cross talk point analysis of a network is a critical approach to understanding how different pathways and modules interact and influence each other. It identifies important points which are interacting with multiple pathways and playing a crucial role in regulating the network. We have calculated the cross-talk point by subtracting the degree of node of the individual pathway from total degrees of node. From cross talk point analysis, we have found acetyl-CoA, malonyl-CoA, propionyl coenzyme A, serine, argininosuccinate, α-KG, tryptophan, carbamoyl aspartate, citrate, valine and glutamine as the important cross talk points with higher non-zero values. Among all of them acetyl-CoA are showing highest interaction with branch chain amino acid metabolism, tryptophan metabolism, TCA cycle, fatty acid synthesis and acetylation pathway (Figure 4).

**Figure 4.**
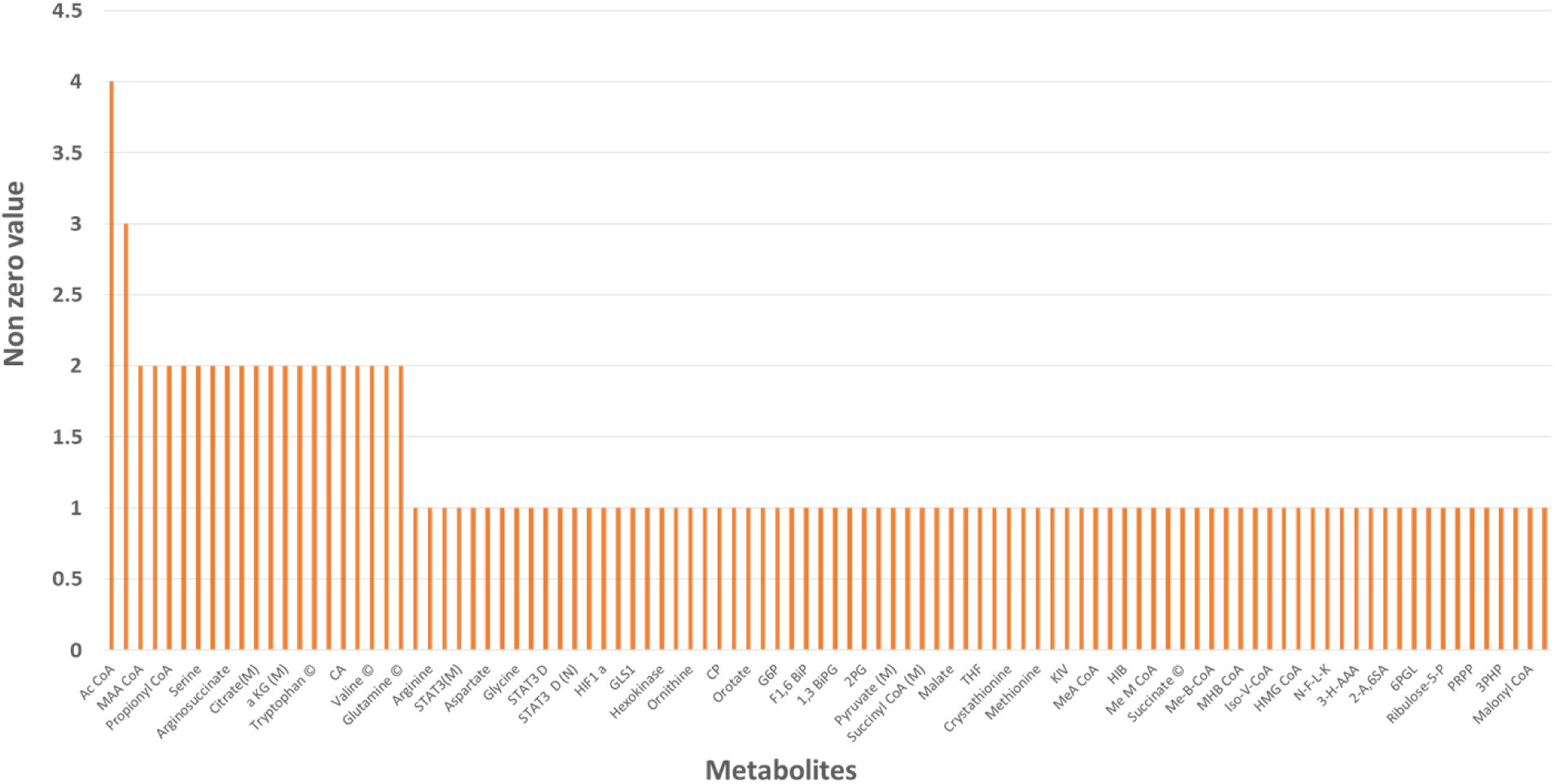
Cross talk points identified from the reconstructed mathematical model having non-zero value greater than or equal to one

### Network analysis

We have constructed a network of our model and used Cytoscape for visualization and analysis of the network (S3). This builds biological networks by representing molecules like genes or proteins as nodes and their interactions as edges, allowing users to create networks and analyses like clustering, hub detection, and functional enrichment. The network was represented in the circular layout and analysed (Figure 5a). The summary statistics of our network show a total 232 species (nodes), 328 connections (edges) and average number of neighbors of a node is 2.7. The number of longest shortest path between any two nodes (diameter) is 22 and shortest path length of a node (radius) is 12. The clustering coefficient value 0.4 (Figure 5b).

**Figure 5.**
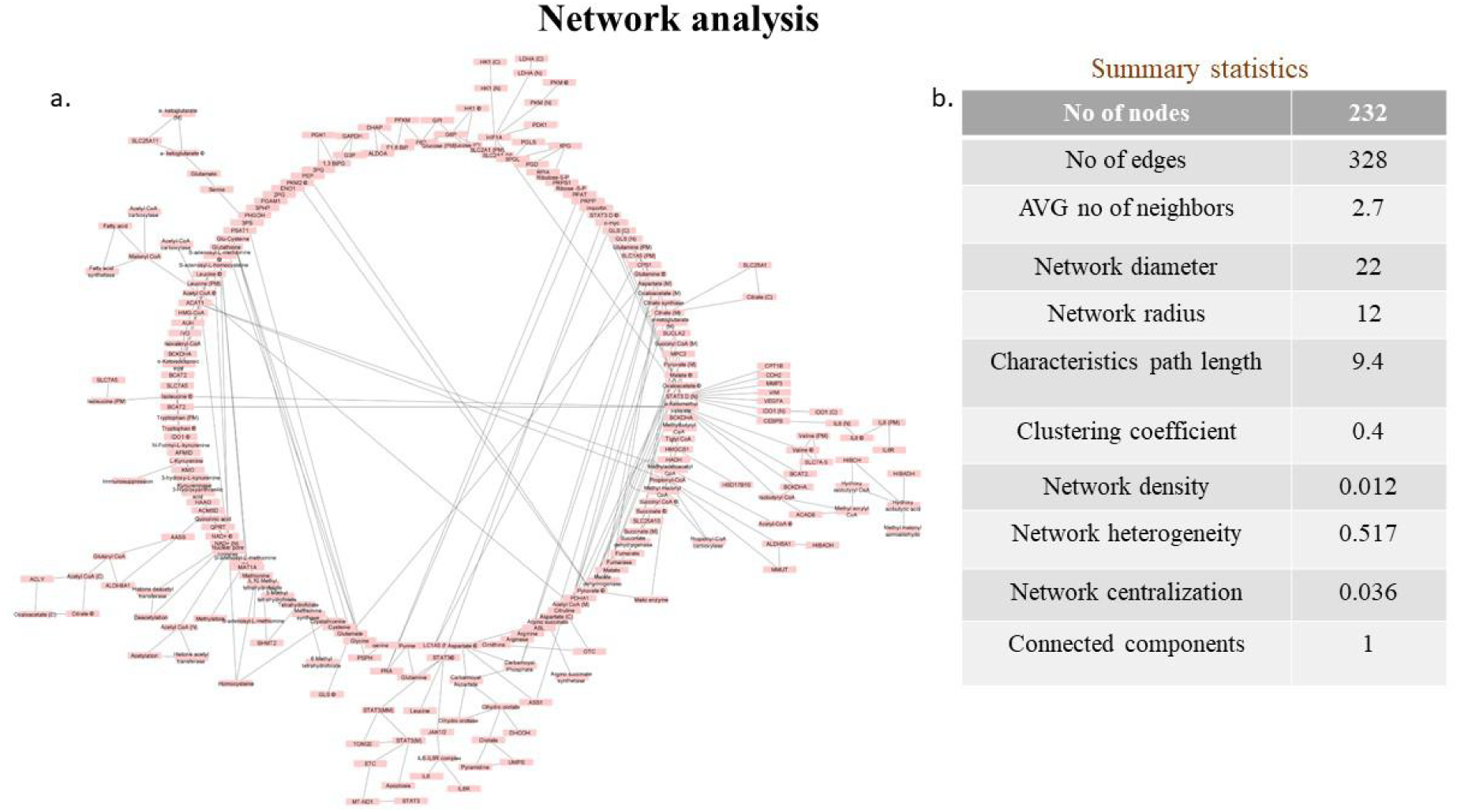
Network construction of the mathematical model (a) Circular layout representation of the mathematical model in the form of network (b) Table showing the parameters of the analyzed network

The network’s top-ranking nodes were identified using a variety of topological analysis techniques using CytoHubba. Total 12 parameters were used to identify key nodes based on MCC, MNC, DMNC, EPC, radiality, degree, proximity, betweenness, eccentricity, bottleneck, clustering coefficient, and stress centrality.

MCC identifies the hub nodes with maximal cliques, a node’s involvement in maximal cliques, indicates a node’s participation in forming stable connections in the network and might be functionally significant (Figure 6a). MNC helps to determine the size of the largest cluster of connected neighbors by examining a node’s immediate neighbors (Figure 6b). The EPC assesses the robustness of a node’s connections under random edge removal, helping to identify nodes critical for maintaining network integrity (Figure 6c). Degree centrality measures the number of direct connections a node has, with high-degree nodes acting as a central node in the biological processes (Figure 6d). DMNC evaluates the density of connections within the MNC, providing insight into how connected a node is within a cluster (Figure 6e). Bottleneck centrality identifies nodes that act as connecting bridges, corresponding to regulatory nodes and acts as signal mediators essential for information flow and network stability (Figure 6f). Betweenness centrality quantifies how often a node appears on the shortest paths between other nodes, identifying key nodes for information flow and communication (Figure 6g). Closeness centrality calculates the inverse of the average shortest path length from a node to all other nodes, identifying nodes that can efficiently communicate across the network (Figure 6h). Radiality measures how close a node is to all other nodes relative to the network’s diameter. High radiality suggesting that the node has a central role in communicating the information flow in the network (Figure 6i). Stress counts the number of shortest paths passing through a node, highlighting those critical for connecting multiple pathways which may possess a regulatory control on biological functions (Figure 6j). Eccentricity represents the greatest distance between two nodes. Nodes of having low eccentricity might be centrally located and may rapidly influence the information flow in the network (Figure 6k). The measure of the degree where nodes tend to cluster together in a network is measured by clustering co-efficient (Figure 6l). The top ten nodes in the network were obtained by analyzing all 12 grading parameters and frequency of occurrence plot was generated. The most important node in our network analysis identified was acetyl-CoA, STAT3 and aspartate (Figure 7).

**Figure 6.**
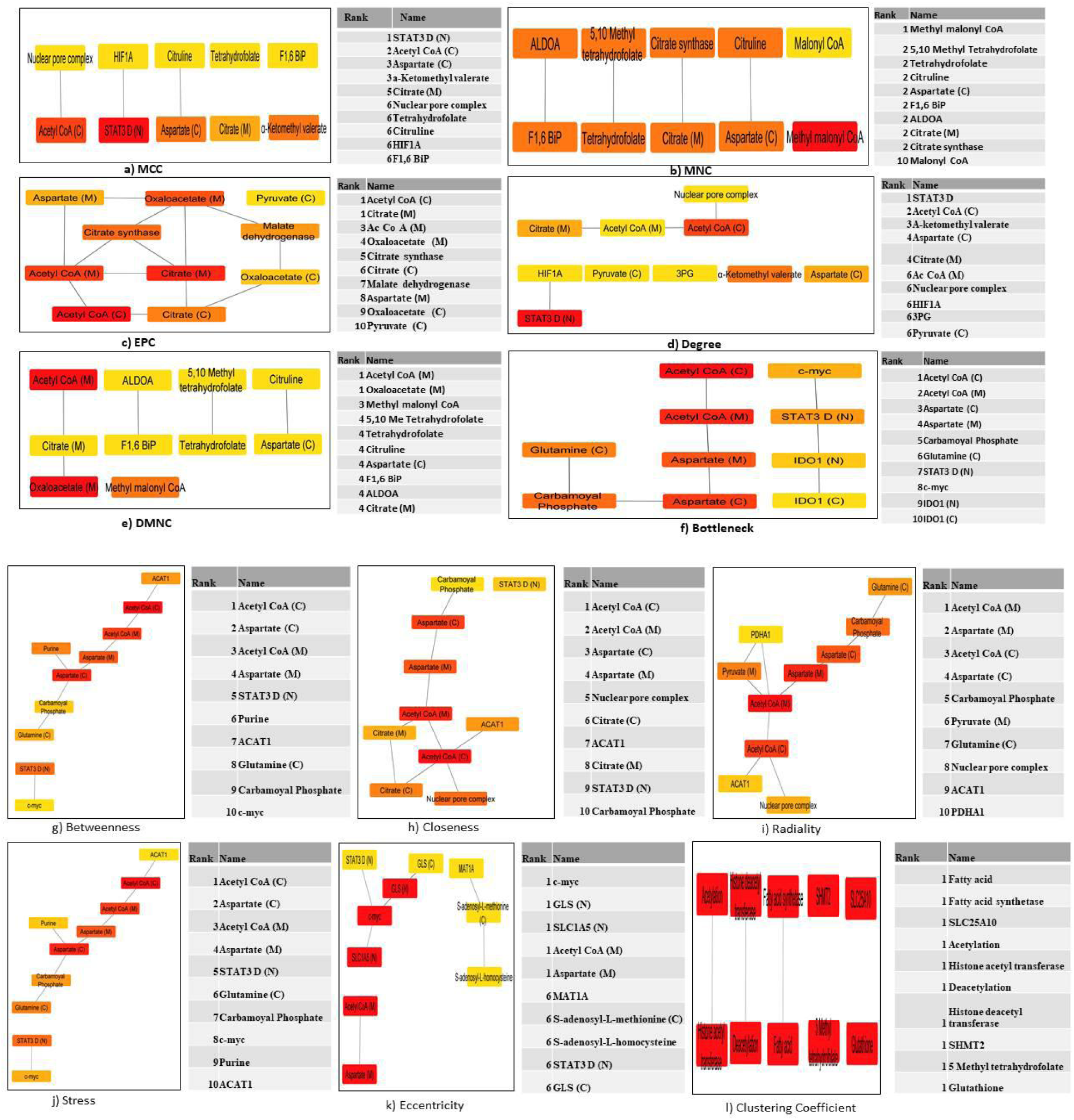
CytoHubba analysis of the network (a)MCC (b)MNC (c)EPC (d)Degree (e)DMNC (f)Bottleneck (g)Betweenness (h) Closeness (i)Radiality (j)Stress (k)Eccentricity (l)Clustering Co-efficient

**Figure 7.**
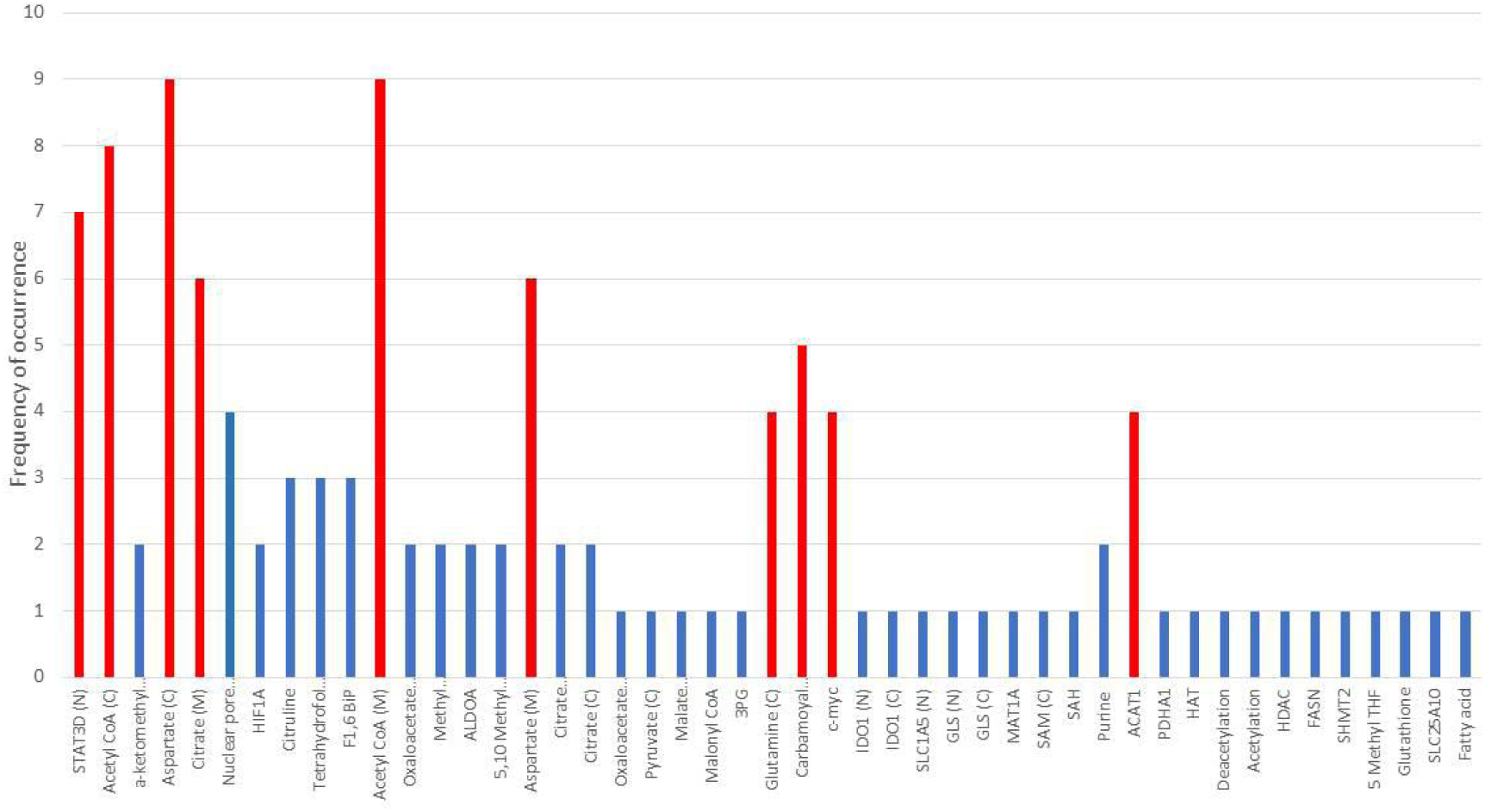
Frequency of occurrence plot of network components evaluated based on CytoHubba parameters. The components highlighted in red are components with frequency of occurrence in 4 or more than 4 parameters. The components highlighted in blue are components with frequency of occurrence less than 4

For network analysis, "BiNGO" (Biological Networks Gene Ontology tool) is used to analyze Gene Ontology (GO). By using this we identified significantly enriched GO categories, providing insights into the functional themes of the studied gene set (Figure 8a) (S4). We found that glucose catabolic process, amine metabolic process, nucleotide metabolic process and anaerobic respiration is significantly enriched in our cancer model (Figure 8b).

**Figure 8.**
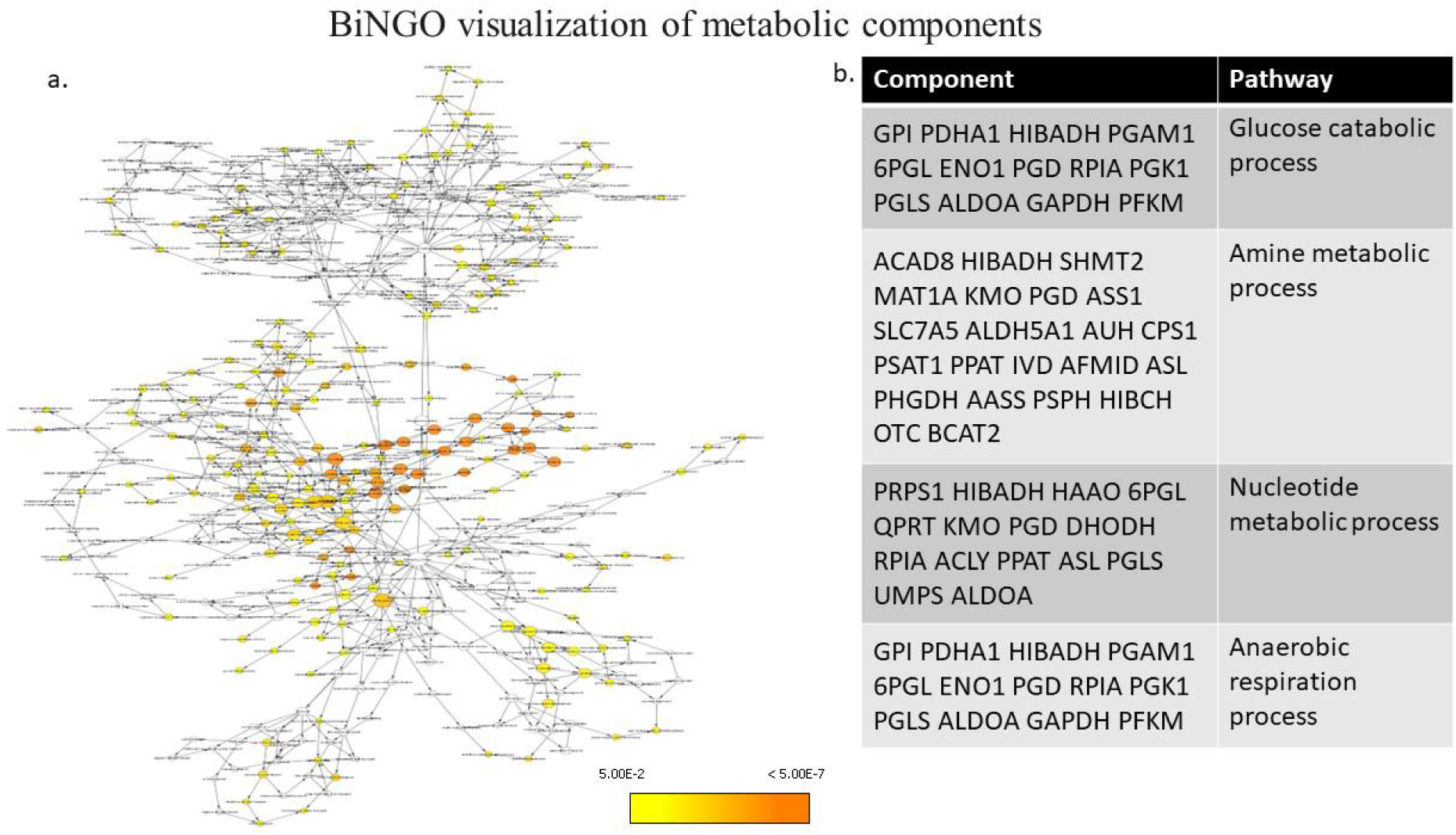
BiNGO analysis of over-represented GO pathways. (a) Graphical representation of GO upon over-representation analysis of the network. (b) Table of components and pathways enriched through BiNGO analysis

Localized cellular expression analysis of Arginase1, IL-6, P-AKT, AKT and STAT3 in H1299, H1975 and A549 upon IL-6 induction was performed. The results show that IL-6, Arginase and AKT expression was significantly upregulated in the H1299 cell line upon treatment with IL-6 (Figure 9a-9f). In the H1975 cell line significant expression change was observed in AKT, P-AKT and STAT3 levels upon IL-6 induction (Figure 9g) (S5). Upon IL-6 induction in the A549 cell line, expression of IL-6, Arginase and AKT expression was significantly upregulated (Figure 9h) (S5). This observation suggests that IL-6 induction leads to the upregulation of IL-6 mediated signaling in NSCLC cell lines.

**Figure 9.**
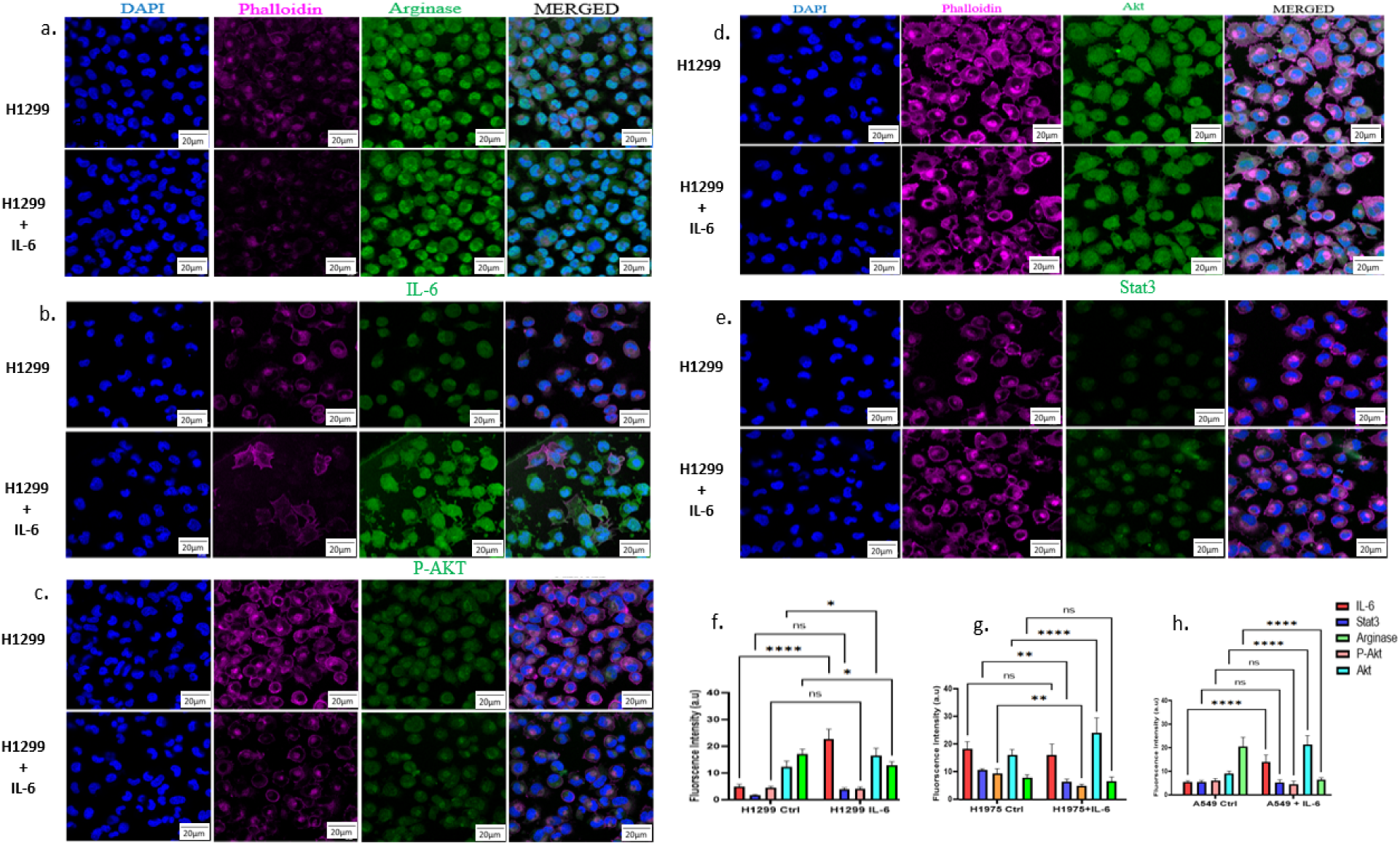
Laser scanning confocal microscopy of H1299 cell line with and without IL-6 induction. The single channel images show DAPI staining (blue), Phalloidin staining (magenta) and antibody staining (green) and merged images of all channels. Antibodies used included-(a) Arginase (b) IL-6 (c) P-AKT (d)AKT and (e) STAT3. Fluorescence intensity was measured for all proteins in cell lines with and without IL-6 induction represented in the graphs (f) H1299 and H1299 induced with IL-6 (g) H1975 and H1975 induced with IL-6 (h) A549 and A549 induced with IL-6. Two-way ANOVA with Dunnet’s multiple comparison test was used for comparing fluorescence intensities. p-value obtained <0.05 was considered significant.

Western blot analysis shows expression change of the proteins upon IL-6 induction in H1299, H1975 and A549 cell lines. We observed significant change in P-STAT3 (Y705) expression in the H1299 cell line induced with IL-6 suggesting active IL-6 signaling in this cell line (Figure 10a and 10b). STAT3 was identified as a cross talk in the model as well as a key regulator in the network. qRT-PCR data shows significant upregulation in ACAC in H1299 cell line induced with IL-6 suggesting upregulated acetyl-CoA expression which we have obtained as a central player of BCAA metabolism (Figure 10c). ELISA for IL-6 expression analysis shows that IL-6 levels in the H1299 cell line are significantly upregulated upon induction whereas, in the H1975 cell line there was no significant difference observed. In A549 we observed downregulation of IL-6 upon induction with IL-6, which was significant (figure 10d). These results suggest that H1299 might be suitable cell line model which may show help in understanding IL-6 signaling role in BCAA metabolism in NSCLC model to understand sarcopenia.

**Figure 10.**
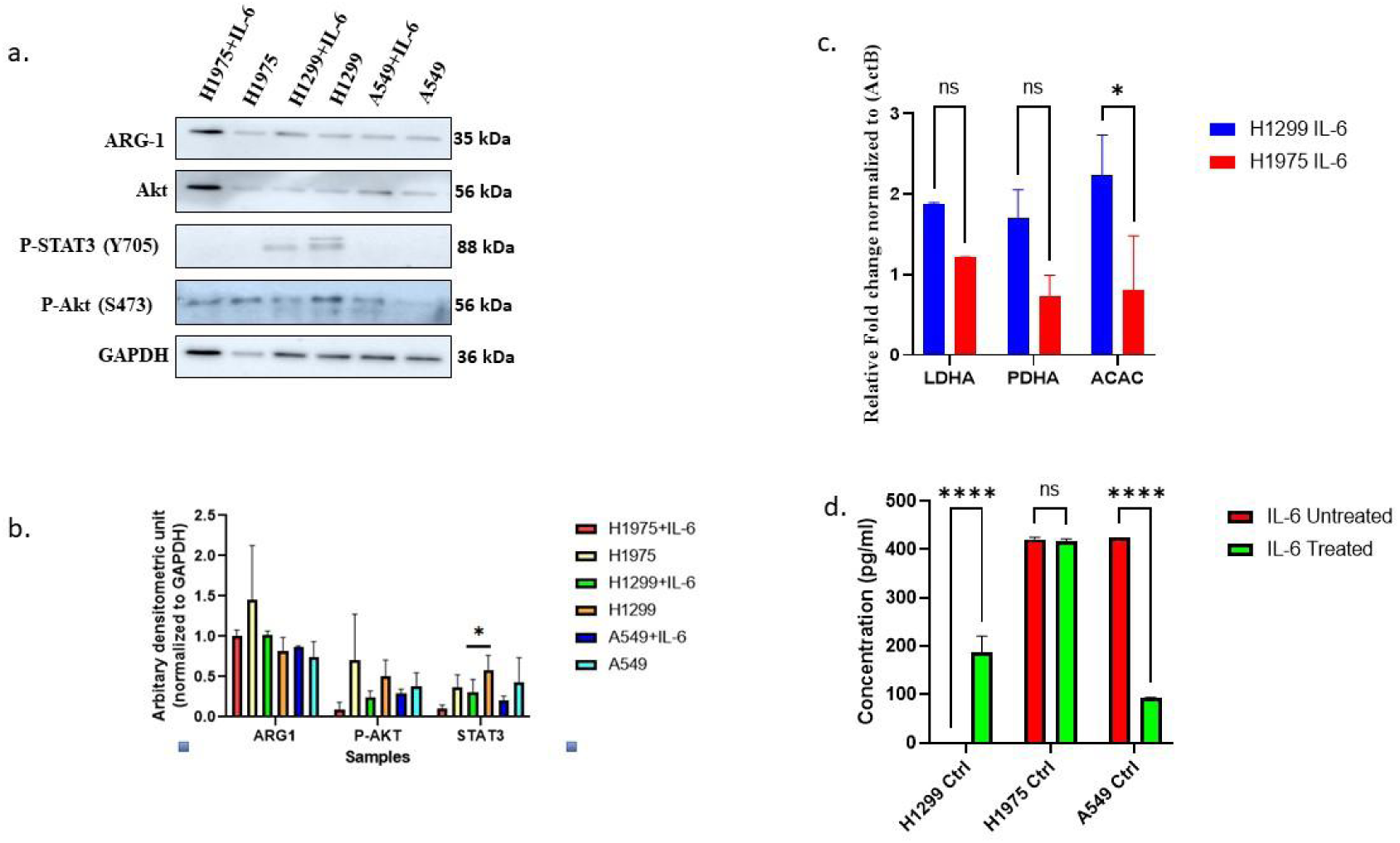
Validation of the mathematical model using NSCLC cell lines- (a) Western blot of NSCLC cell line with and without IL-6 induction for expression analysis of Arginase, AKT, P- AKT, STAT3 and GAPDH. (b) Densitometry analysis of western blot normalized with β-actin. Two-way ANOVA with Bonferroni’s multiple comparison test was used for comparing expression of proteins. p-value obtained <0.05 was considered significant. (c) qRT-PCR analysis of LDHA, PDHA and ACAC. Two-way ANOVA with Bonferroni’s multiple comparison test was used for comparing expression fold change. p-value obtained <0.05 was considered significant. (d) ELISA for IL-6 expression. Two-way ANOVA with Bonferroni’s multiple comparison test was used for comparing expression of IL-6. p-value obtained <0.05 was considered significant.

## Discussion

The reconstructed mathematical model illustrating dysregulated immune-metabolism in NSCLC cells shows the interdependence of IL-6 signaling pathway regulating nutrition through BCAA and redox potential through mitochondria. Both these pathways are essential and are prime reasons for inducing sarcopenia in patients suffering from NSCLC. The representative model highlights the role of crucial hallmarks of cancer which include avoiding immune destruction, tumor-promoting inflammation, activating invasion and metastasis, inducing vasculature, and epigenetic reprogramming which leads to cancer progression and may potentially induce sarcopenia. The IL-6 mediated JAK-STAT3 signaling induces genes involved in cancer cell metabolic pathways which trigger glycolysis, amino acid metabolism, redox homeostasis, and fatty acid metabolism.

It also triggers biological processes that induce metastasis and angiogenesis. In the model dual role of STAT3 phosphorylation states have been presented to play different roles. STAT3 Y705 phosphorylation translocates STAT3 to the nucleus whereas STAT3 S727 phosphorylation translocates STAT3 to mitochondria to regulate ETC and inhibit the intrinsic pathway of apoptosis. Interactive simulation of the model was performed using SOSlib solver to determine concentrations of the model species based on the kinetic laws used in the model against time course [40]. From the simulation analysis of the model, we observed fatty acid metabolism, acetylation reactions, glutathione synthesis, pyrimidine synthesis, deacetylation reactions and methylation reactions increasing with time. Species which showed reduced production yet prominent concentrations included acetyl-CoA, succinate, malonyl-CoA and NAD+. The initiation and progression of NSCLC is due to a constant interplay between robust and permanent epigenetic alterations [41]. These alterations primarily occur at DNA and histones which may result in DNA methylation, histone acetylation and deacetylation [42]. Increased expression and improved uptake of BCAAs have been linked to elevated glutamate and glutamine levels in NSCLC tumors. Nitrogen produced by BCAA deamination aids in the production of nucleotides and non-essential amino acids in NSCLC [43]. The end products of BCAA oxidation differ depending on the specific amino acid, as studies have demonstrated that only the catabolism of valine and isoleucine generates carbon substrates capable of supporting gluconeogenesis independently of acetyl-CoA [44]. A crucial metabolic intermediate, acetyl-CoA is engaged in several important physiological functions, such as the TCA cycle, the synthesis of fatty acids, and protein acetylation that plays a role in epigenetic control. acetyl-CoA helps regulate tumor growth, aggressiveness, and drug resistance by linking histone acetylation to lipid metabolism. The high reactivity of its thioester bond, which supports pathways like glycolysis, the TCA cycle, de novo lipogenesis, and the mevalonate pathway, is crucial to its metabolic processes. Despite not being membrane-permeable, acetyl-CoA can diffuse between the cytosol and nucleus and operates independently in peroxisomes, mitochondria, and the cytosol [45]. Thus acetyl-CoA acts as key player in controlling the metabolism of NSCLC.

PCA is a versatile exploratory method that is ideal for evaluating a variety of numerical data types since it does not rely on distributional assumptions [46]. With its wide capability, the R package FactoMineR is especially used for multivariate data analysis. It allows for the incorporation of variables from structural information in the data, and supports both quantitative and categorical variable types. Additionally, the dimensions that emerge from exploratory studies can also be automatically described by means of category and/or quantitative variables [47]. All the species from the model were categorized into two groups: Species which were a part of a single pathway and species which participated directly in multiple pathways. The variable plot suggested that species from both categories have PCA score more than 1. The species with PCA score more than 1 were identified from the individual plot and represented by their pathways in the PCA plot. Species contributing to high PCA score was identified by percentage contribution.

Thus highlighting, FASN, MDH, succinyl-CoA synthetase, Importin, Glutamate-cysteine, Arginase and Methylation as PCs from individual pathways. 5,10 methyl THF, acetyl-CoA (M), acetyl-CoA (N), NAD+, Dihydroorotase (DHOase), Dihydro orotate and IL-6 as PCs from multiple pathways. PHP, LDHA, Pyruvate Dehydrogenase, α-KG and Glutamate as PCs which may be crucial for all pathway functioning.

Flux analysis revealed that high flux reactions are promoting production of succinate, acetyl-CoA, NAD+, SAM, α-KG, citrate, aspartate, pyruvate, isoleucine, valine, leucine and STAT3. The high flux reaction analysis draws an insight into the relationship between IL-6 signaling with metabolism of BCAA which might be regulated by Krebs cycle reaction intermediates; more specifically acetyl-Coa. Cross talk point analysis identified acetyl-CoA, malonyl-CoA, propionyl-CoA, Serine, Argininosuccinate, citrate, α-KG, tryptophan, carbamoyl aspartate (CA), Valine and Glutamine with cross talk non zero value more than 1. Among all the cross-talk points STAT3 stood out as the cross-talk point which is a part of IL-6 immune signaling; rest other cross talk points were enzymes and metabolites participating in metabolic pathways.

The network analysis’s objective was to pinpoint significant components in the network by investigating their functional role and connectedness [48]. The clustering coefficient of the network was 0.4 suggesting moderate connectivity within the nodes of the network. CytoHubba analysis and analyzing the frequency of occurrence of nodes in top 10 network analysis parameters suggests that STAT3, acetyl-CoA, aspartate, citrate, nucleopore complex, glutamine, Carbamoyal phosphate, c-Myc and acetyl-CoA acetyltransferase (ACAT1). CytoHubba results suggest that the enlisted components are crucial based on the interconnectedness of the model as a network for regulation of immune-metabolic pathways. BiNGO analysis identified glucose catabolic process, amino acid metabolism, nucleotide metabolic process and anaerobic respiration pathways as overrepresented pathways.

The present study provides comprehensive insights into the differential activation of IL-6-mediated signaling pathways across NSCLC cell lines; H1299, H1975, and A549 upon IL-6 induction, highlighting the cell-type-specific nuances that may underlie their contribution to sarcopenia-related mechanisms through BCAA metabolism. Our localized cellular expression analysis and Western blot data collectively demonstrate that H1299 cells exhibit robust upregulation of IL-6, Arginase1, AKT, and P-STAT3 (Y705), indicating a highly active IL-6 signaling axis. This is further supported by qRT-PCR data revealing significant upregulation of ACAC, a key regulator in acetyl-CoA production and BCAA catabolism, thereby implicating IL-6-driven metabolic rewiring in this cell line. Conversely, H1975 cells primarily show altered expression of AKT, P-AKT, and STAT3, suggesting partial pathway activation, while A549 cells exhibit upregulation of IL-6, Arginase, and AKT, yet a paradoxical downregulation of IL-6 protein levels by ELISA, implying a complex feedback regulation. These discrepancies underscore the heterogeneity of IL-6 responsiveness among NSCLC subtypes, with H1299 emerging as a particularly responsive model that may best mimic the IL-6–BCAA metabolic axis potentially contributing to muscle wasting in cancer. Our findings support the role of STAT3 and acetyl-CoA as a central regulatory node and potential cross-talk mediator in this network, reinforcing the utility of H1299 cells as a relevant *in vitro* system to further elucidate the molecular links between chronic inflammation, cancer metabolism, and sarcopenia.

The IL-6–BCAA immuno-metabolism mathematical model represents a cutting-edge approach for understanding and addressing NSCLC-induced sarcopenia. By integrating key metabolic and inflammatory pathways, the model highlights the pivotal interplay between STAT3 signaling and acetyl-CoA metabolism, both of which are central to muscle wasting in the context of lung cancer. This systems-level framework not only advances mechanistic insight into tumor-driven metabolic dysregulation but also identifies actionable nodes within the network—such as STAT3 activation and acetyl-CoA flux—that can be strategically targeted using precision medicine to mitigate sarcopenia in NSCLC patients.

### CRediT authorship contribution statement

Gautam Kumar: Writing – review & editing, Writing –original draft, Visualization, Validation, Software, Resources, Methodology, Investigation, Formal analysis, Data curation.

Shweta Khandibharad: Writing – review & editing, Writing –original draft, Visualization, Validation, Software, Resources, Methodology, Investigation, Formal analysis, Data curation. Shailza Singh: Writing – review & editing, Supervision, Software, Resources, Project administration, Methodology, Investigation, Funding acquisition, Formal analysis, Conceptualization.

## Acknowledgements

We thank Department of Biotechnology, Ministry of Science and Technology, Government of India for intramural funding. We also thank Director, Biotechnology Research and Innovation Council-National Centre for Cell Science (BRIC-NCCS), Pune for supporting the Bioinformatics and High-Performance Computing Facility at BRIC-NCCS.

